# Amino Sugar Metabolism with AgaSY Coordinates Carbon Catabolism, Fimbrial Regulation, and Pathogenesis in Uropathogenic *E. coli*

**DOI:** 10.64898/2026.10.16.759770

**Authors:** Carole Anamalé, Hicham Bessaiah, Sébastien Houle, Sima Mohammadi, Roger C. Levesque, Charles M. Dozois

## Abstract

Uropathogenic *Escherichia coli* (UPEC) exploit alternative carbon sources in order to adapt to the nutrient-limited urinary tract. While glucose is often scarce in healthy human urine, UPEC express a variety of sugar transport and catabolic systems that promote fitness during infection. In this study, we identify the *agaSY* genes in UPEC strain CFT073 as essential for the catabolism of the amino sugars N-acetyl-galactosamine (GalNAc or Aga) and galactosamine (GalN or Gam) and for the efficient use of additional amino sugars and carbon sources. Deletion of *agaSY* not only abolished Aga utilization but also impaired catabolism of multiple carbon sources, including amino acids and sugars tested using phenotypic arrays. The Δ*agaSY* mutant exhibited significantly reduced type 1 fimbriae production, reduced biofilm formation, and increased motility, suggesting a link between amino sugar metabolism and the regulation of bacterial surface structures. Loss of *agaSY* did not affect growth and adherence to bladder cells and increased expression of genes encoding F9 fimbriae. Furthermore, the Δ*agaSY* mutant was significantly attenuated for bladder and kidney colonization in a murine urinary tract infection model, while complementation restored urinary tract colonization. The results indicate that a combination of deficiencies in nutrient acquisition, altered carbon metabolism, and regulation of virulence-associated factors contributes to reduced urinary tract colonization. Our findings highlight the interplay between metabolic flexibility and virulence regulation, as well as the central role that amino sugar metabolism plays in UPEC pathogenesis.

**Author Summary:** Urinary tract infections are among the most common bacterial infections, and most are caused by uropathogenic *Escherichia coli.* To survive and cause infection, these bacteria need to take up nutrients from the urinary tract, where the available nutrients can be limited. In this study, we were interested in the impact of the ability of *E. coli* to utilize different nutrients and to use them to cause urinary tract infections. We focused on two genes that allow *E. coli* to metabolize amino sugars, a group of nutrients that can be found in the urinary tract. We found that these genes are important not just for using these sugars but also for using many other nutrients. Knocking out these genes affected several mechanisms such as surface structure production, motility and biofilm formation. Most importantly, bacteria lacking these genes were less able to colonize the bladder and kidneys in mice, and restoring the genes restored this ability. We demonstrate a close link between nutrient use and the regulation of virulence factors, particularly fimbrial genes. Understanding this link may help to clarify how *E. coli* can adapt to the urinary tract and identify new ways to interfere with bacterial infection.

## Introduction

Metabolic pathways are closely linked to the virulence of pathogenic bacteria, making them attractive targets for antibacterial strategies. In uropathogenic *Escherichia coli* (UPEC), several key metabolic circuits have been implicated in its ability to colonize and persist in the urinary tract. Urinary tract infections (UTIs) rank among the most common infectious diseases worldwide, affecting hundreds of millions of individuals each year (1, 2). Women are particularly vulnerable to UTIs, largely due to anatomical differences in the urogenital tract. Epidemiological data indicate that over 50% of women will experience at least one UTI during their lifetime, with a substantial proportion experiencing recurrent infections (3). Furthermore, 20-30% of women who experience an initial UTI will develop a recurrent infection within six months, underscoring the urgent need to better understand the underlying microbial mechanisms of UTI pathogenesis (4).

In approximately 80% of community-acquired UTIs, the causative agent is UPEC (1, 5). As a highly adaptive pathogen, UPEC can evade host defenses, persist within the urinary tract, and may harbor genes to withstand antibiotic treatment. The bladder constitutes a nutrient-limited niche, driving UPEC to evolve metabolic strategies that not only ensure survival but also enable proliferation under these restrictive conditions. These bacteria exhibit remarkable metabolic strategies, allowing them to utilize a range of carbon sources present in urine at low concentrations, including sugars, peptides, amino acids, and host-derived metabolites. This metabolic flexibility is increasingly recognized as a virulence trait that supports both survival and proliferation under restrictive conditions (6). However, merely characterizing virulence factors does not provide a comprehensive understanding of infectious disease mechanisms. It is also important to investigate how pathogenic bacteria modulate their metabolic pathways to exploit carbon and energy sources within the host environment, thereby promoting their growth, persistence and successful colonization at sites of infection. The emergence of drug-resistant UPEC strains further compounds the clinical challenge, representing a significant threat to global health. Therefore, understanding the mechanisms underlying UPEC metabolism may provide new avenues for potential treatment and prevention of UTIs.

Metabolic pathways play a pivotal role in UPEC adaptation to these challenges. For example, in nutrient-limited urine, the *gat* operon enables utilization of galactitol, a sugar alcohol derived from galactose, thereby supporting bacterial growth (7). In the oxygen-limited environment of the bladder, the transcriptional regulator FNR governs the expression of anaerobic respiration genes (8), while siderophore systems facilitate ferric iron uptake (Fe^3+^), an essential micronutrient that is of limited availability in urine (5, 9). Moreover, gluconeogenesis and tricarboxylic acid (TCA) cycle pathways have been shown to contribute to UPEC fitness in the murine urinary tract (7). Collectively, these pathways illustrate how metabolic adaptations complement classical virulence strategies to promote infection.

Another critical aspect of UPEC virulence is the vast array of virulence factors that facilitate colonization and survival in the urinary tract. UPEC adhere to and invade the bladder through surface structures, including lipopolysaccharide (LPS), polysaccharide capsules, flagella, outer membrane vesicles and proteins, as well as secreted toxins, siderophores and fimbrial adhesins. Among these, type 1 fimbriae (pili) play a predominant role in virulence. Encoded by the *fim* operon, these adhesins play a critical role in the early stages of infections caused by UPEC (5, 10). Recently, our laboratory investigated the role of novel regulators in the expression of type 1 fimbriae. Using a transposon (Tn) library in UPEC reference strain CFT073, we identified several mutants exhibiting both reduced stress tolerance and altered type 1 fimbriae production. Notably, multiple genes involved in the metabolism of N-acetyl-D-galactosamine (GalNAc or Aga) and D-galactosamine (GalN or Gam) were identified to affect *fim* expression (11, 12). In addition, the *kbaZ-agaV and kbaY-agaSBC* operons, which mediate the transport and metabolism of Aga and Gam, respectively, were also found to be down-regulated in a UPEC CFT073 mutant lacking the *ryfA* gene, which encodes a small RNA (sRNA)(13).

Amino sugars, such as glucosamine (GlcN) and N-acetyl-glucosamine (GalNAc or Nag), play an important role in the synthesis of the structural components of Gram-negative bacteria, including peptidoglycan and lipopolysaccharides (LPS)(14, 15). In many bacteria, the transport and concomitant phosphorylation of these carbohydrates are mediated by substrate-specific, membrane-bound Enzyme II (EII) complexes of the phosphoenolpyruvate (PEP)-dependent carbohydrate phosphotransferase system (PTS). Specifically, the amino sugars Aga and Gam are transported into the cell via dedicated PTS systems encoded by the 11.7 kb *aga*/*gam* gene cluster. This gene cluster comprises two operons and codes for two EII complexes and for enzymes involved in the catabolism of these sugars. For Gam, the *agaB*, *agaC*, and *agaD* genes encode the three subunits of the AgaBCD (EII^Gam^) complex. Similarly, for Aga, the *agaV*, *agaW*, *agaE*, and *agaF* genes form the AgaVWEF (EII^Aga^) complex, responsible for Aga import and phosphorylation (14, 16, 17).

In the present study, using UPEC strain CFT073, we investigated the importance of Aga/Gam sugar metabolism with a particular focus on the *agaSY* genes, which encode an isomerase (*agaS*) and aldolase (*agaY*) in the Aga/Gam metabolic pathway. Deletions of the *agaSY* genes generated in UPEC CFT073 were found to play a broader role in carbon metabolism and also resulted in virulence attenuation in a mouse UTI model. Our results suggest a bifunctional role for *agaSY* genes in the Aga/Gam pathway, as these genes were shown to be required not only for catabolism of Aga and Gam but also contributed to the metabolism of additional sugars and influenced expression of virulence-associated traits, such as regulation of fimbriae.

## Results

### Deletion of *agaSY* genes markedly affects carbon catabolism in *E. coli* CFT073

The GalNAc utilization pathway has been previously reconstructed through comparative genomics in a wide range of *Proteobacteria* species(30) and experimentally characterized in *E. coli*(14), *Streptococcus suis* and *Lactobacillus*(31). Since the utilization of Aga and Gam in UPEC strains is not well studied, we investigated their role in the UPEC reference strain CFT073. Based on prior findings in *E. coli* and *in silico* analyses, we hypothesized that the Aga/Gam catabolic pathways in UPEC strain CFT073 resemble those of reference strain *E. coli* C (Figure 1)(14).

**Figure 1.**
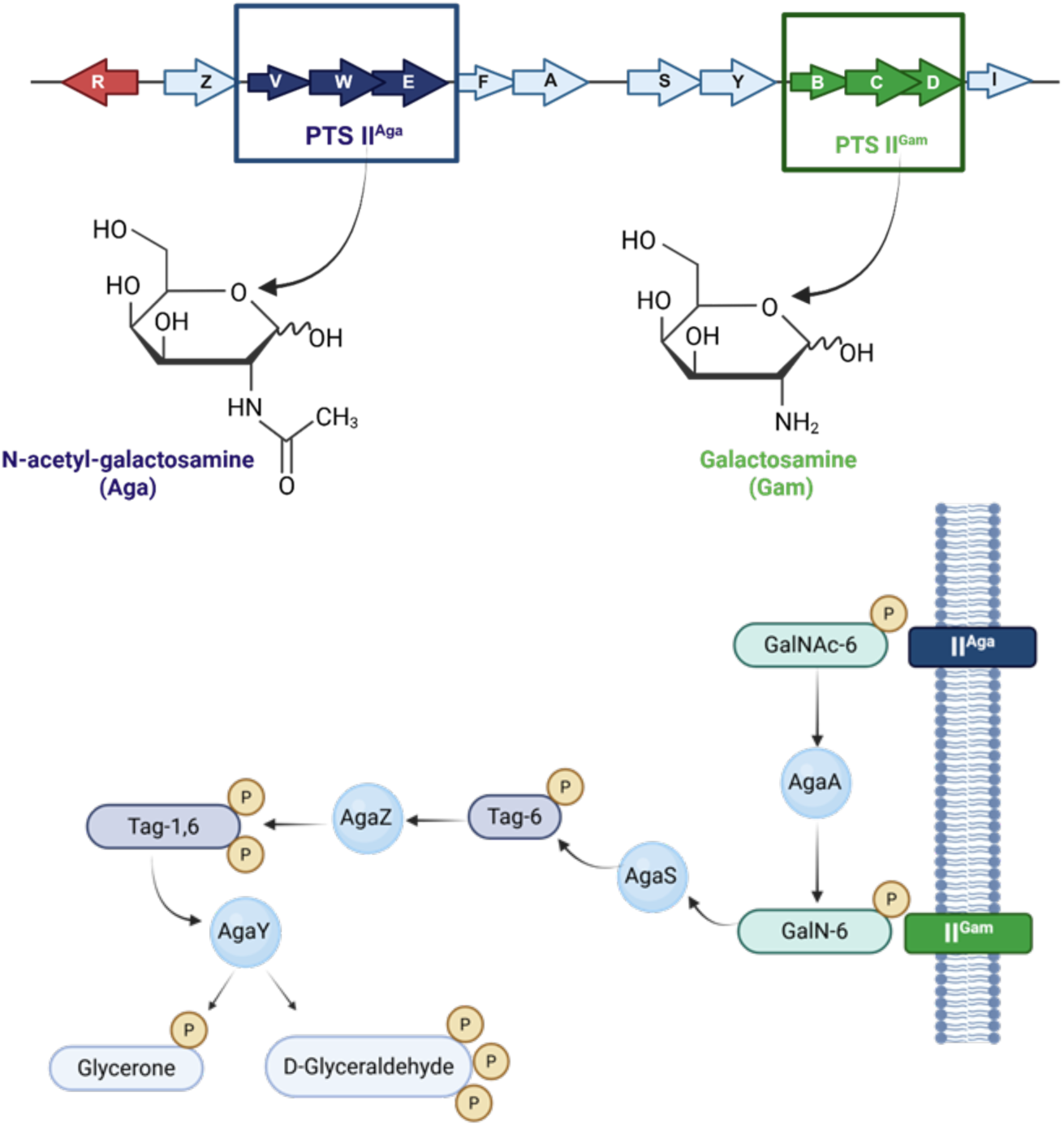
Organization of the *aga* gene cluster and catabolic pathway of Aga/Gam transport and utilization in *E. coli*. In *E. coli*, Aga and Gam are transported and phosphorylated by their respective PTS system (EII^Aga^ and EII^Gam^) to GalNAc-6-P and GalN-6-P. GalNAc-6-P is deacetylated by AgaA to GalN-6-P, which is then converted by AgaS to tagatose-6-P, phosphorylated by AgaZ to tagatose-1,6-biphosphate, and finally cleaved by AgaY into glycerone and D-glyceraldehyde.

As both GalNAc (Aga) and GalN (Gam) have been shown to serve as efficient carbon sources for some *E. coli* strains, which contain the full set of *aga* genes, we investigated the role of *agaSY* in their utilization. To evaluate the role of *agaSY* in this process, we constructed a Δ*agaSY* mutant in UPEC strain CFT073 and assessed whether *agaSY* are required for catabolism of these sugars or other carbon sources. CFT073, its derivative Δ*agaSY* mutant, and the complemented strain, as well as *E. coli* K-12 and *E. coli* W strains, were evaluated for their ability to utilize Aga or Gam as the sole carbon source. After 48 hours of incubation at 37°C, the results demonstrated that *agaSY* are essential for Aga catabolism in the CFT073 wild-type strain. Consistent with previous findings(14)*E. coli* K-12, which carries a deletion of the *agaVWEFA* cluster, was unable to grow on either Aga or Gam and thus served as a negative control. In contrast, *E. coli* W, similar to *E. coli* C, exhibited rapid growth on both sugars and was used as a positive control.

Importantly, the Δ*agaS*Y mutant did not grow on Aga-containing minimal medium, whereas growth was restored in the complemented strain. Although the CFT073 wild-type strain contains the complete *aga*/*gam* gene cluster, including the *agaBCDI* genes required for Gam transport, it was unable to grow on Gam as the sole carbon source, and the deletion of *agaSY* did not influence this phenotype (Table 1, Fig.2A). To compare growth on solid versus liquid media, we performed kinetic growth assays for at least 70 hours at 37°C. Consistent with the observations on agar plates, the CFT073 wild-type strain and *E. coli* W exhibited a similar growth rate on Aga (Fig. 2C), whereas the Δ*agaSY* mutant displayed markedly reduced growth, comparable to that of the *E. coli* K-12 strain. *E. coli* W was the only strain capable of growing in M9 supplemented with Gam (Fig. 2B and 2D). All strains grew at comparable rates in LB or M9 supplemented with glucose (Fig.S1).

**Figure 2.**
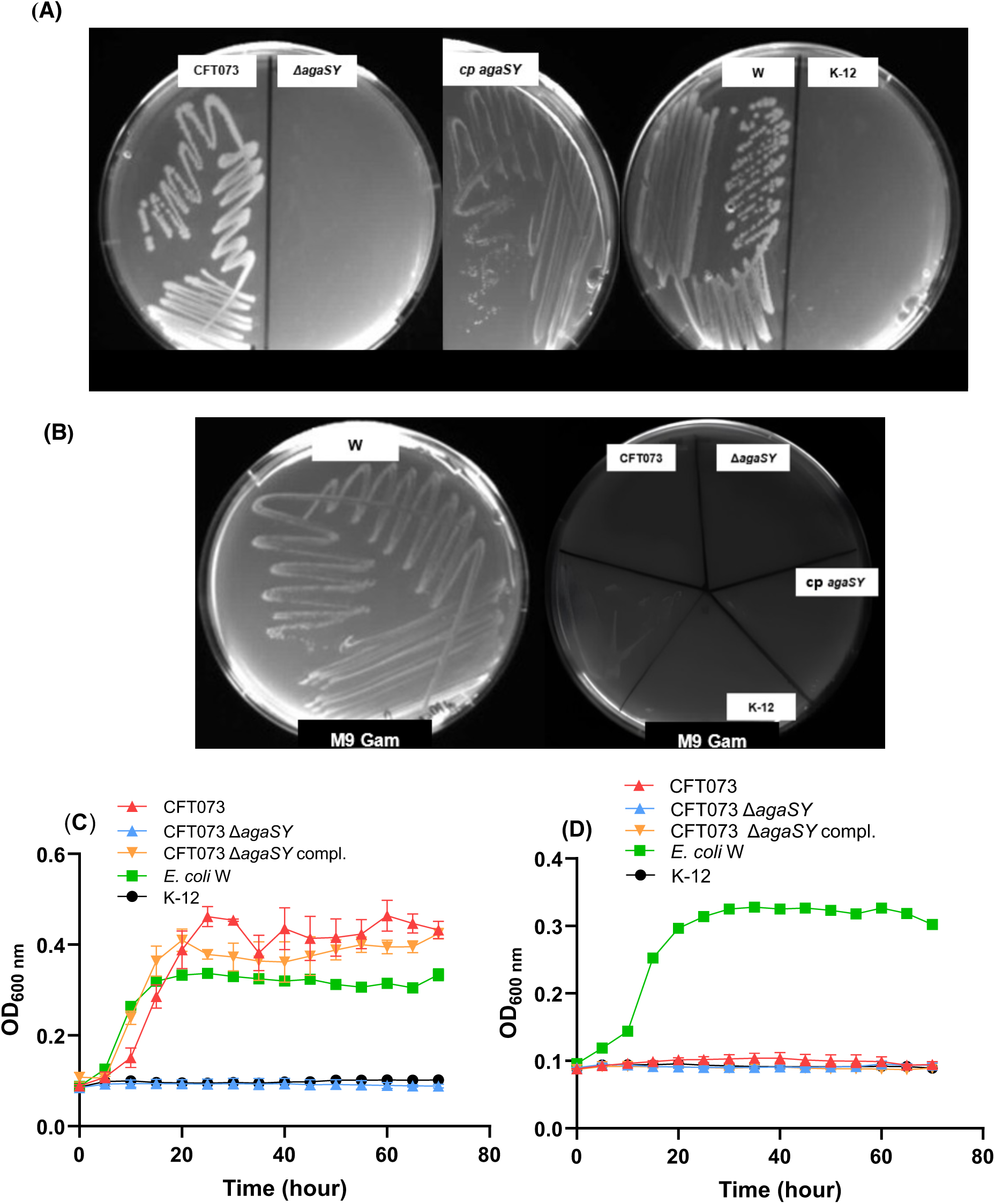
Growth and kinetic growth of CFT073 wild-type and Δ*agaSY* mutant on minimal M9 medium supplemented with (<u>A, C</u>) Aga and (<u>B, D</u>) Gam. Growth of CFT073 UPEC strain and its derivative Δ*agaSY* mutant and complemented strain on M9 0.2% Aga agar plates (A); lack of growth occurred with 0.2% Gam on plates (B); Growth in M9 liquid 0.2% Aga (C), and in M9 liquid 0.2% Gam (D). For growth on plates, strains were first grown overnight (16-18 hours) on LB agar at 37°C. A single colony of each strain was re-isolated onto M9 supplemented with 0.2% Aga (A) or 0.2% Gam (B). For kinetic growth, strains were grown overnight (16-18 hours) in LB broth at 37°C with shaking, washed with PBS, and inoculated into fresh M9 medium supplemented with 0.2% Aga (C) or 0.2% Gam (D). Cultures were incubated at 37°C for 70 hours, and growth was monitored. In both panels, the red line represents CFT073 wild-type strain, the blue line the Δ*agaSY* mutant, the orange line the complemented *agaSY* strain, the green line *E. coli* W (positive control) and the black line *E. coli* K-12 (negative control).

**Table 1.** Bacterial growth in M9 medium supplemented with 0.2% Aga or 0.2% Gam.

| Strains | Growth with Aga | Growth with Gam |
| --- | --- | --- |
| *E. coli* K-12 | - | - |
| *E. coli* W | + | + |
| CFT073 | + | - |
| CFT073 Δ*agaSY* | - | - |
| CFT073 Δ*agaSY* compl. | + | - |

### Deletion of *agaSY* genes reduces biofilm and type 1 fimbriae and increases motility

To investigate the role of *agaS,* we first constructed a chromosomal deletion in the UPEC CFT073 strain. Given the well-established importance of type 1 fimbriae in UPEC adhesion, we assessed the effect of the *agaS* deletion on type 1 fimbriae production using the yeast agglutination assay (Figure 3A). When cultures were grown in LB at 37°C to mid-log phase (OD_600_ ≈ 0.6), the Δ*agaS* mutant displayed a significant decrease (*p*<0.01) in yeast agglutination compared to the CFT073 wild-type strain. Complementation of *agaS* alone only partially restored agglutination, as the titers remained lower than those of the wild-type parental strain (Fig.3A). Examination of the genomic region of *agaS* within the *aga* gene cluster suggested that complementation of *agaS* alone was insufficient, as downstream genes of *agaS* may also be required. Indeed, a promoter, p*agaS*, immediately upstream of *agaS* has been identified in *E. coli*, suggesting the possibility of co-regulation (14). To test this hypothesis, we generated a Δ*agaY* mutant and observed the same decrease in agglutination as in the Δ*agaS* mutant. Further, complementation with *agaS* and *agaY* together fully restored the wild-type phenotype (Fig. 3A). Based on these results and considering the genetic organization of the *agaSY* genes, we used a Δ*agaSY* double mutant for further characterization in this study. These results suggest that loss of *agaSY* caused a substantial decrease in the production of type 1 fimbriae.

**Figure 3.**
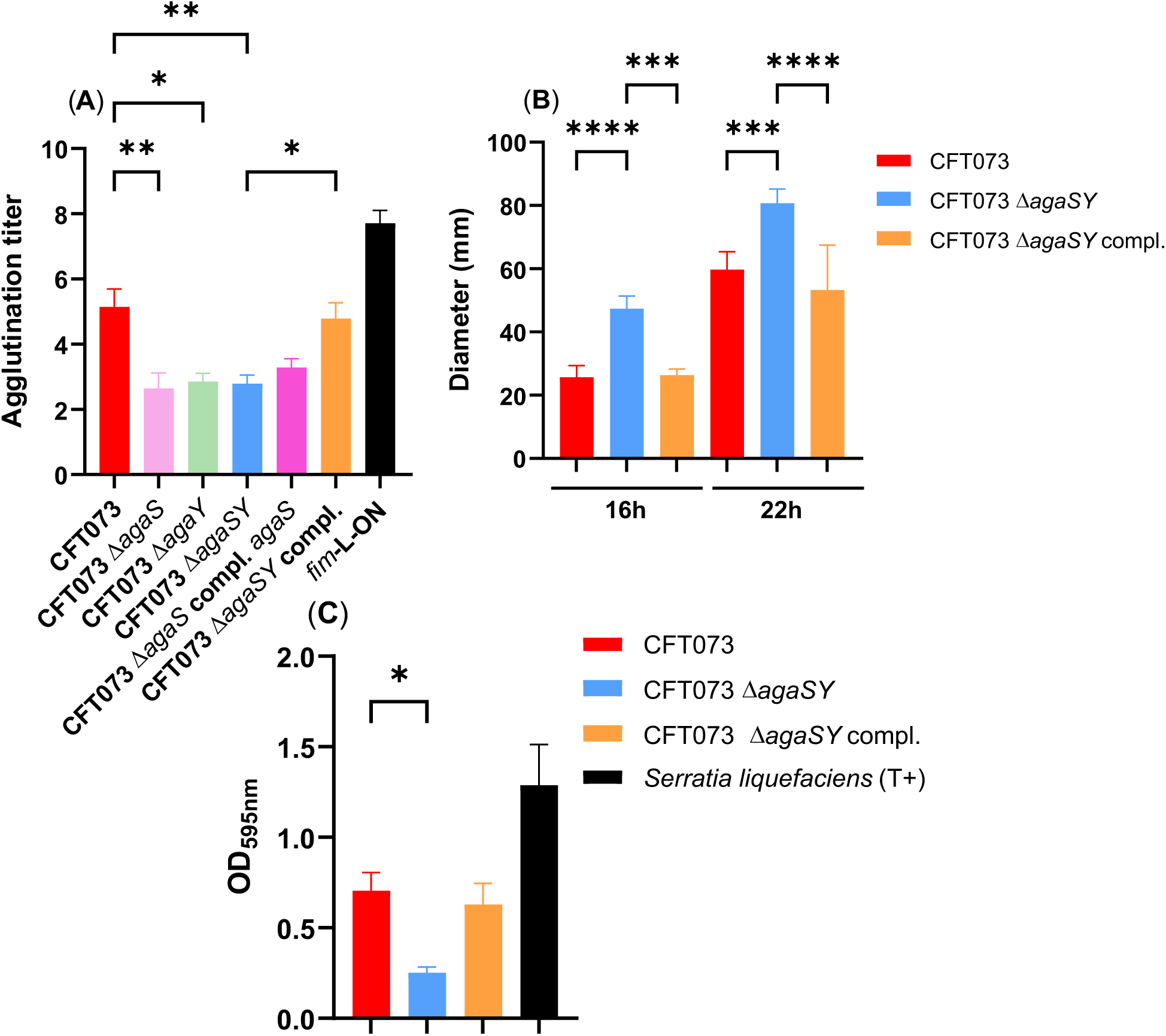
Effect of deletion of *agaSY* genes for type 1 fimbriae production, motility and biofilm formation. (**A**) Yeast agglutination titer of CFT073 wild-type strain and its derivative mutants *agaS, agaY, agaSY* and the complemented *agaS* and *agaSY* strains demonstrating type 1 fimbriae production in strains cultured to mid-log phase in LB medium. The CFT073 *fim*-locked ON strain was used as a positive control. (**B**) Motility of CFT073, the Δ*agaSY* mutant and the complemented strain on 0.25% soft agar. Bars represent the mean motility zone diameter, with error bars indicating standard deviations from six independent biological replicates. (**C**) Biofilm formation assay using the crystal violet assay for CFT073 wild-type, Δ*agaSY* mutant, the complemented *agaSY* strain and the positive control *Serratia liquefaciens.* All statistical significance was calculated by the Kruskal-Wallis test, followed by Dunn’s multiple comparisons test; * *p≤* 0.05; ** *p≤* 0.01; *** *p≤* 0.001; **** *p≤* 0.0001.

Since effects on the *fim* operon can influence the flagellar system by increasing or decreasing bacterial motility, and as motility contributes to host colonization and persistence (26, 32), we also performed a motility assay on 0.25% soft agar plates. Deletion of *agaSY* resulted in a significant increase (*p*<0.0001) in motility after 16 and 22 hours of incubation at 37°C (Fig. 3B).

As biofilm formation is a critical step in UTI, enabling UPEC to proliferate, accumulate and persist within the host by evading immune defenses(1, 5), we also examined the effect of *agaSY* deletion on biofilm formation in UPEC CFT073. The biofilm formation ability was quantified using a crystal violet assay. Cultures were grown in LB medium to mid-log phase and incubated in a plastic 96-well plate at 37°C for 48 hours to allow biofilm formation. The Δ*agaSY* mutant displayed a significant reduction in biofilm density compared to the wild-type CFT073 strain (*p*<0.05). Complementation with *agaSY* restored biofilm formation to levels comparable to those observed with the CFT073 wild-type strain (Figure 3C). Taken together, these findings highlight the critical role of *agaSY* genes and their effects on motility, as well as the regulation of type 1 fimbriae and biofilm formation.

### Deletion of *agaSY* genes significantly reduced murine urinary tract colonization

Since nutrients in urine are limited, consisting mainly of dilute amino acids and small peptides, bacteria must utilize available resources to persist in the urinary tract. Previous studies have shown that defects in type 1 fimbrial expression or production reduce UPEC virulence *in vivo* (13, 33, 34). Consistent with these findings, using a murine UTI model, we hypothesized that loss of *agaSY* would impair urinary tract colonization. Infection experiments were conducted with 10 mice per group, each inoculated with the CFT073 wild-type strain, the *agaSY* mutant or the complemented strain. Bacterial colonization was assessed in the bladder and kidneys 48 hours post-infection (pi). The Δ*agaSY* mutant exhibited significantly reduced bacterial loads in both kidneys (Figure 4A, right and left Figure 4B) and the bladder (Figure 4C) compared with the wild-type strain (*p*<0.01 and *p*<0.05). As expected, complementation of *agaSY* resulted in a regain of bacterial burden to levels that were not significantly different from the wild-type (Figure 4). Collectively, these results suggest that the enzymes encoded by *agaSY*, known to contribute to catabolism of Aga and Gam, are also important for colonization of the bladder and kidneys by UPEC CFT073 in a murine UTI model.

**Figure 4.**
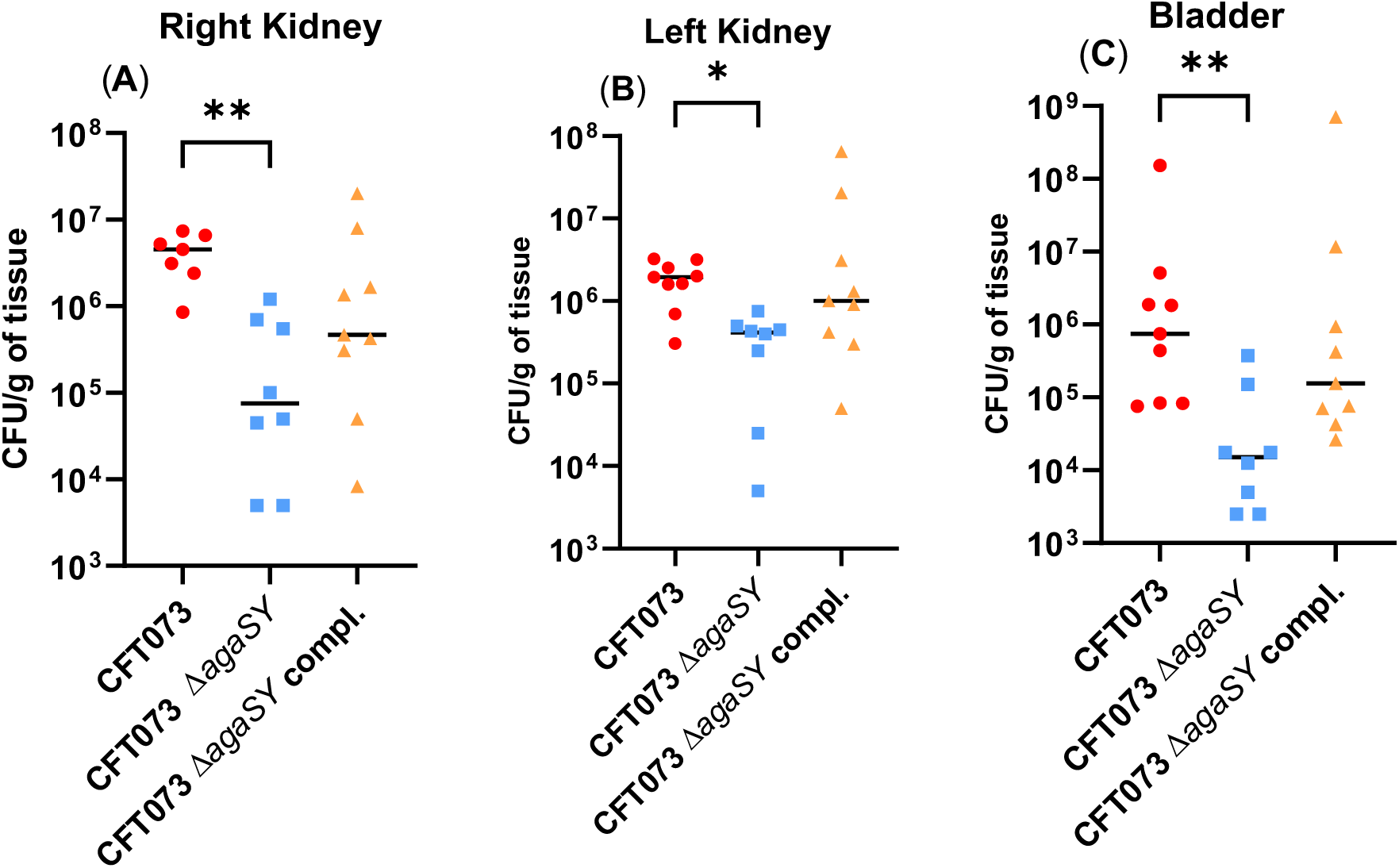
Lack of *agaSY* induces bacterial virulence attenuation in a murine UTI model. Single-strain infections were performed with 10 mice per group to compare CFT073 wild-type strain, the Δ*agaSY* mutant and the Δ*agaSY* complemented strain. Inocula were prepared from cultures grown in BHI medium, and 1.20 to 2.40 × 10^9^ CFU/30µL were administered per mouse. Bacterial loads were quantified in (**A**) right kidney, (**B**) left kidney, and (**C**) bladder at 48 hours post-infection (pi). Each data point represents an individual mouse, and horizontal bars indicate median values. Statistical significance was assessed using the Kruskal-Wallis test, a non-parametric alternative to one-way ANOVA, followed by Dunn’s post-hoc multiple comparisons test;* *p ≤* 0.05; ** *p ≤* 0.01.

### Deletion of *agaSY* affects expression of genes encoding other fimbrial adhesins

As amino sugars such as Aga or Gam are common constituents of bacterial cell wall surface structures(14, 17), we hypothesized that changes in carbon metabolism due to deletion of *agaSY,* which resulted in reduced *fim* expression and increased motility, could potentially affect expression of other fimbrial systems. Since UPEC strains adhere to bladder epithelial cells primarily through type 1 fimbriae(25, 35), we reasoned that the *in vivo* attenuation of the CFT073 Δ*agaSY* mutant could, at least in part, be attributed to impaired type 1 fimbriae-mediated colonization of the bladder. To test this hypothesis, we compared the adherence capacity of strains to 5637 bladder cells. As shown in Fig. 5A, in the absence of 2.5% α-D-mannopyranose, the adherence of the *agaSY* mutant was comparable to that of the wild-type and complemented strains. The CFT073 Δ*fim* strain, used as a control, also displayed similar adherence to the wild-type strain (Fig. 5A). Since type 1 fimbriae recognize mannosylated residues on host epithelial cells, the addition of soluble mannose to the culture medium should inhibit type 1 fimbriae-dependent adhesion. Interestingly, upon addition of 2.5% α-D-mannopyranose, the Δ*agaSY* mutant exhibited significantly higher bacterial adherence in comparison with the wild-type strain (Fig. 5A). Therefore, this unexpected phenotype suggested that in the *agaSY* mutant, other adhesins or fimbriae could potentially compensate for the decreased production of type 1 fimbriae.

**Figure 5.**
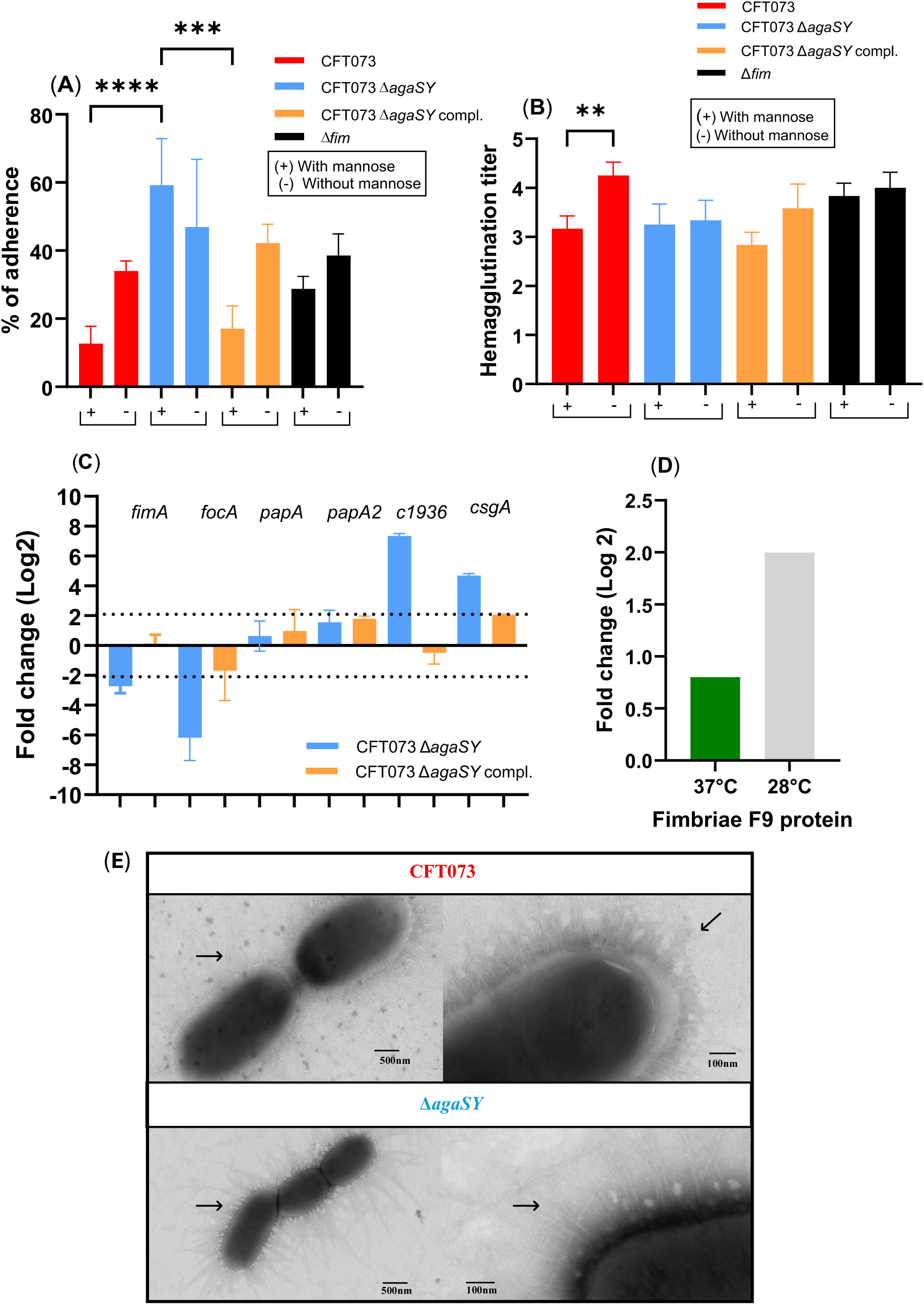
Implication of *agaSY* in the regulation of fimbrial adhesin production. (**A**) Adherence of strain CFT073 and its derivatives mutant Δ*agaSY* to human 5637 bladder epithelial cells in the presence or absence of 2.5% α-d-mannopyranose was determined. (**B**) Production of Pap fimbriae by hemagglutination assay. Results are the mean values and standard deviations for three biological experiments. Statistical significance was assessed using the Kruskal-Wallis test, a non-parametric alternative to one-way ANOVA, followed by Dunn’s post-hoc multiple comparisons test; * *p ≤* 0.05; ** *p ≤* 0.01;*** *p ≤* 0.001; **** *p ≤* 0.0001. (**C**) Expression of fimbrial genes*, fimA*, *focA*, *papA*, c1936, and *csgA* by qRT-PCR in the Δ*agaSY* strain compared to CFT073 wild-type strains. Cultures were grown in LB until mid-log exponential phase. (**D**) Quantification of F9 fimbrial proteins by LC-MS/MS. The relative abundance of F9 fimbriae in CFT073 Δ*agaSY* compared to the CFT073 wild-type strain is expressed as fold change (log2) at 37°C and 28°C. (**E**) Electron microscopy of CFT073 and the Δ*agaSY* mutant. Arrows show fimbriae on cell surfaces.

In UPEC strain CFT073, previous studies have shown that repression of type 1 fimbriae can lead to increased expression of Pap fimbrial genes expression (36, 37). To assess whether Pap fimbriae were upregulated in the Δ*agaSY* mutant, we performed mannose-resistant hemagglutination (MRHA) assays using human erythrocytes. As shown in Fig. 5B, no significant difference in hemagglutination was observed between CFT073 wild-type and the Δ*agaSY* mutant, suggesting that Pap fimbriae production was not markedly altered.

To further investigate the potential contribution of other adhesins, we quantified the expression of some other fimbrial genes present in strain CFT073, including *papA* and *papA_2* (Pap fimbriae), *focA* (F1C fimbriae) and c1936 (F9 fimbriae). Surprisingly, deletion of *agaSY* results in substantial differences in fimbrial gene expression. Expression of *focA* (F1C) was down-regulated 64-fold, whereas expression of *csgA* (curli) was up-regulated around 16-fold and expression of F9 fimbriae(38) was up-regulated 256-fold. Additionally, *fimA* expression was reduced 8-fold, as evidenced by the decrease observed in type 1 fimbriae-associated yeast agglutination observed in the Δ*agaSY* mutant (Fig. 5C). Expression of genes encoding Pap fimbriae was not significantly altered. To confirm whether this transcriptional upregulation of F9 fimbriae translated into increased protein production, we performed LC-MS/MS quantification of F9 fimbrial proteins. As shown in Fig. 5D, the CFT073 Δ*agaSY* mutant exhibited an increased abundance of F9 fimbrial proteins compared to the wild-type strain, particularly at 28°C. These results are consistent with regulatory changes in the expression of multiple fimbrial systems in UPEC CFT073, due to the loss of the *agaSY* genes, and suggest that F9 fimbriae could potentially compensate for the decrease in type 1 fimbriae observed.

Further, consistent with the adherence phenotype to 5637 bladder epithelial cells (Fig. 5A) and the potential for increased production of F9 fimbriae (Fig. 5C and Fig. 5D), electron microscopy images revealed morphological differences at the bacterial surface, with distinct fimbrial structures observed in the Δ*agaSY* mutant compared to those produced by the CFT073 wild-type strain (Fig. 5E).

These findings support the hypothesis that the loss of type 1 fimbriae in the Δ*agaSY* mutant is at least partially compensated for by the production of other fimbriae or adhesins. Moreover, the strong upregulation of other fimbriae in the *agaSY* mutant indicates altered regulation of fimbrial adhesins due to disruption of Aga/Gam catabolism, indicating a potential link between altered carbon metabolism and modified regulation of adhesins produced by UPEC.

### Phenotypic Microarray

Phenotypic Microarrays (PMs) from Biolog^TM^ can be used to directly assay the effects of genetic changes on cells, especially gene knockouts. To assess whether the *agaSY* genes contribute to the metabolism of sugars and other carbon sources other than Aga and Gam, the wild-type CFT073 strain and its isogenic Δ*agaSY* mutant were tested for carbon utilization assays using the PM1 plate, which contains a total of 96 carbon sources, including specific amino acids. Following the manufacturer’s protocols, we performed a comparative phenotypic analysis of strain CFT073 and the Δ*agaSY* mutant. The PM1 assay provided an overview of their physiological characteristics (Figure 6), revealing that CFT073 metabolized multiple carbon sources efficiently. By contrast, the deletion of *agaSY* resulted in a substantial impairment in selective carbon source utilization compared to the parental strain (Figure 6). Interestingly, the Δ*agaSY* mutant was unable to grow on N-acetylglucosamine (Aga), one of the most abundant natural sugars that serves as both a carbon and nitrogen source (A3) or when provided with mannose as the sole carbon source (A11) and with L-alanyl-glycine (G6), L-glutamine (E1) or L-alanine (A9), which represent the dominant amino acids naturally present in human urine (39).

**Figure 6.**
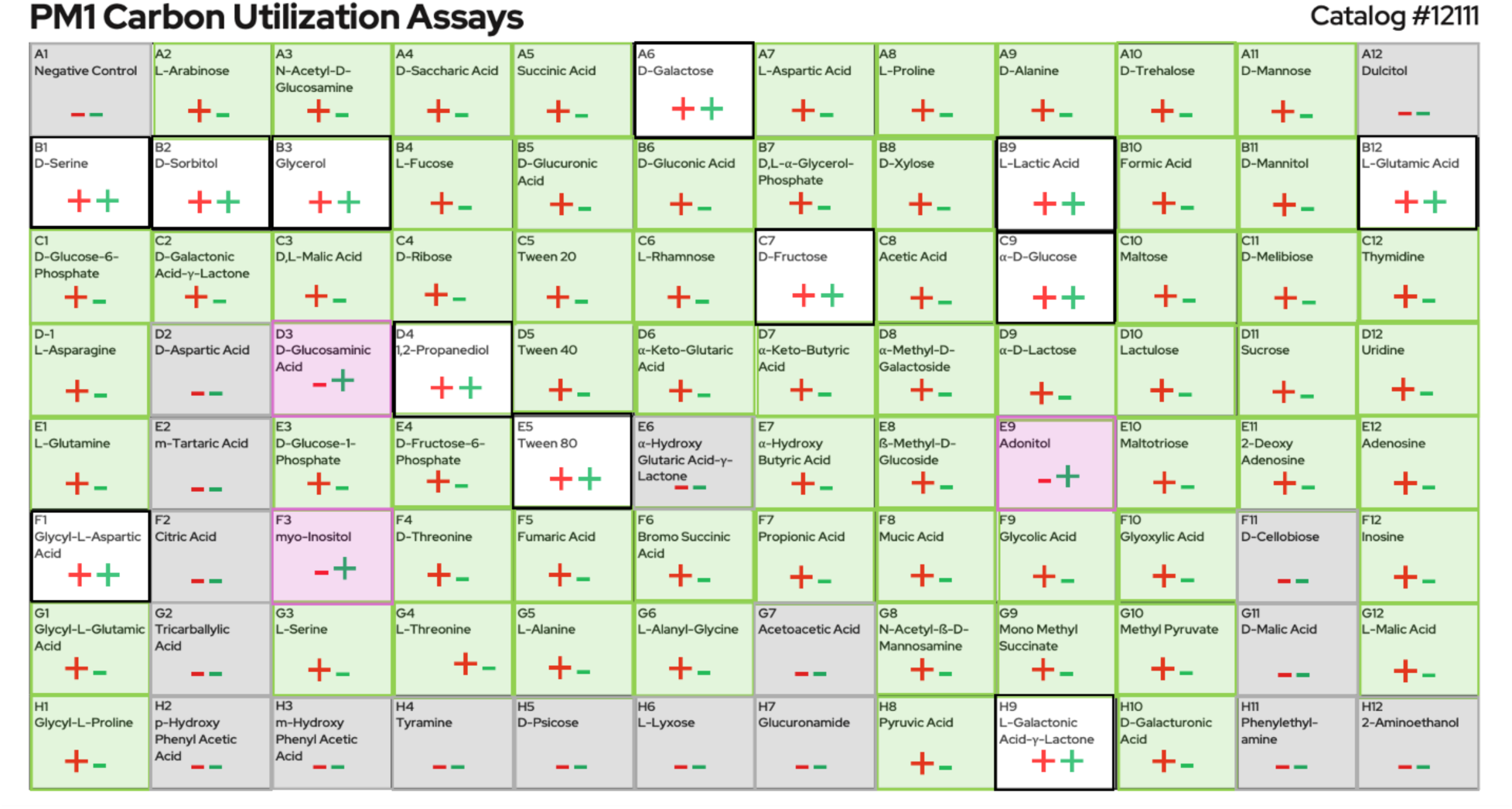
Comparisons of growth obtained from CFT073 wild-type and the Δ*agaSY* mutant screened on PM1 (Carbon utilization) Each well of the 96-well PM1 (A01-H12) plate represents a distinct substrate and displays growth/metabolic activity of the wild-type strain, CFT073, indicated by red (+/-) symbols for positive or negative utilization of the corresponding carbon source and the mutant Δ*agaSY,* indicated by green (+/-) symbols. Background colors indicate differential growth; grey wells show no growth by either strain; green wells indicate growth by the wild-type strain only; white wells indicate growth by both strains, and pink wells indicate growth by the mutant strain only.

Together, these results demonstrate that in UPEC CFT073, *agaSY* is not only required for Aga or Gam utilization but that these genes may also contribute to broader metabolism of carbon sources, including some specific sugars, amino acids, and other compounds.

## Discussion

Bacterial adaptation to nutrient-limiting host environments, such as the bladder, for UPEC is largely dependent on the utilization of carbohydrates, as evidenced by the wide range of sugar transport systems present in *E. coli*. While glucose is the preferred carbon source, this sugar is frequently only available at limited concentrations that prevent growth. To overcome this restriction, *E. coli* has developed strictly controlled secondary nutrient acquisition mechanisms to maintain competitiveness and survival. Here, we demonstrated that the *agaSY* genes in UPEC strain CFT073 are required for the catabolism of Aga and for the effective use of numerous other amino sugars and carbon sources. Further, loss of the *agaSY* genes has pleiotropic effects on important virulence phenotypes, such as production of type 1 fimbriae, motility, biofilm formation and bladder and kidney colonization during urinary tract infection in mice. These findings establish an important role for amino sugar metabolism in the pathogenesis of UPEC and suggest that metabolic genes can contribute to extraintestinal virulence by influencing bacterial physiology beyond nutrient acquisition.

### Impact of *agaSY* deletion on type 1 fimbriae

One of the most important virulence factors for UPEC infection are type 1 fimbriae. Indeed, studies on UPEC CFT073 and UTI89 have already demonstrated that loss or decrease in type 1 fimbriae leads to attenuation in UTI models in mice (10, 40, 41). Adherence to the urinary tract epithelium is the initial stage of a UTI, allowing bacteria to resist the hydrodynamic forces of urine flow and establish an infection. Thus, for successful colonization of hosts by bacteria, it is crucial that the expression of bacterial surface structures, which mediate interaction with host tissues, is finely regulated. In UPEC, multiple and distinct types of regulators have been shown to regulate the production of type 1 fimbriae (42).

Type 1 fimbriae are among the most important adhesins for UPEC, mediating attachment to mannosylated receptors on bladder epithelial cells and initiating colonization and invasion of epithelial cells (25, 43, 44). Previously, several studies have shown that the expression of virulence factors can be modulated by nutrient availability (45); however, the regulatory mechanisms connecting specific metabolic pathways to the regulation of virulence factors remain poorly defined. Within the urinary tract environment, bacteria frequently adapt their metabolism by modulating and regulating their catabolic pathways. In our study, deletion of *agaSY* resulted in a reduced level of type 1 fimbriae production, as demonstrated by a marked decrease in yeast agglutination titer and reduced *fimA* expression by qRT-PCR. Complementation with *agaSY* restored type 1 fimbrial production, confirming the contribution of these genes to the regulation of type 1 fimbriae.

The mechanistic link between amino sugar metabolism and fimbrial regulation remains unclear, but several regulatory pathways may explain the phenotypes observed in the Δ*agaSY* mutant. Carbon source availability is known to influence the expression of type 1 fimbriae through global metabolic regulators, including the CRP-cAMP complex, which controls the expression of the fimbrial recombinases FimB and FimE (46). Disruption of amino sugar catabolism could therefore alter intracellular carbon flux and signaling, leading to changes in the activity of metabolic regulators that ultimately affect fimbrial gene expression. An additional possibility is that perturbation of amino sugar metabolism influences the balance between motile and sessile lifestyles. In *E. coli,* this transition is tightly controlled by the second messenger cyclic-di-GMP, which promotes biofilm formation and adhesive functions while repressing flagellar motility (47). Although our data do not directly demonstrate altered c-di-GMP signaling, metabolic perturbations have been shown to affect c-di-GMP regulatory networks through changes in the activity or expression of diguanylate cyclases and phosphodiesterases. Consistent with this model, the increased motility observed in the Δ*agaSY* mutant may affect a shift toward a more motile state, accompanied by reduced expression of type 1 fimbriae and biofilm formation. The precise molecular mechanism and connection between AgaSY-dependent carbohydrate metabolism and virulence gene regulation remain to be determined, but our findings suggest that disruption of this pathway may influence the balance between fimbrial expression, motility and surface-associated lifestyles through broader regulatory networks.

Further, amino sugar metabolism may influence the envelope stress response, which is known to modulate fimbrial assembly (28, 42). Links between metabolism and virulence have been documented in other contexts; for instance, the presence of the *deoK* locus involved in fructooligosaccharide and deoxyribose metabolism enhances the competitiveness of pathogenic *E. coli* to colonize the murine intestine (48). Both PTS and non-PTS sugar transporters have been involved in extraintestinal pathogenic *E. coli* (ExPEC) pathogenesis, notably with the *gimA* locus, which contributes to host cell infection and survival within host tissues (49). Changes in carbohydrate uptake pathways may therefore indirectly influence fimbrial expression through changes in cellular metabolic status and regulatory signaling pathways that converge on the *fim* switch.

Together, our findings suggest that *agaSY* deletion may similarly disrupt these metabolic or stress pathways, thereby leading to reduced levels of type 1 fimbrial production.

### Effect of *agaSY* deletion on biofilm formation

Biofilm formation represents a critical step in UTI pathogenesis, which allows UPEC to persist within the host, evade immune defenses and resist antibiotic treatment. By proliferating and accumulating on biotic surfaces, UPEC develop biofilms that facilitate colonization of the urinary tract but also allow protection against the immune system, which leads to chronic and recurrent infections (1, 5). Conversely, the transition between planktonic and biofilm-associated lifestyles is tightly regulated, notably through intracellular c-di-GMP signaling, with reduced c-di-GMP levels generally promoting motility at the expense of biofilm formation (47). Given the increased swimming motility observed with the *agaSY* mutant, AgaSY may influence this regulatory network, although this hypothesis remains to be investigated. By mediating attachment to both biotic and abiotic surfaces in the early stages of biofilm formation, type 1 fimbriae play a critical role in this process (50, 51). In our investigation, we observed that deletion of *agaSY* significantly reduced biofilm density *in vitro*, whereas complementation fully restored wild-type biofilm levels.

Several mechanisms may explain this phenotype. First, reduced type 1 fimbrial expression observed in the Δ*agaSY* mutant likely impairs initial surface adhesion. Second, deficient utilization of amino sugars may alter the composition of the cell envelope, which may impact the production of extracellular matrix and surface hydrophobicity. Consistent with these hypotheses, previous studies have shown that biofilm-associated genes, including curli fiber production and cellulose biosynthesis, are regulated in response to nutrient conditions (52, 53). Thus, loss of *agaSY* may indirectly affect biofilm development by altering metabolic flux through these pathways.

### Altered regulation of expression of F9 fimbriae

UPEC genomes encode multiple fimbrial operons, but typically only one type of fimbria is expressed at a time, controlled by regulatory crosstalk and phase variation (54, 55). Inactivation or constitutive expression of a single fimbrial system can modulate the expression of other adhesins (36), enabling bacteria to target diverse host receptors while avoiding immune system detection. For instance, studies have demonstrated that deletion of the *fim* operon may enhance expression of alternative fimbriae (36, 37).

We demonstrated that the deletion of *agaSY* decreased levels of type 1 fimbriae, but did not affect Pap fimbriae production, as indicated by comparable hemagglutination activity and similar expression levels of *papA* and *papA2* genes in both the wild-type and mutant strains. Interestingly, by qRT-PCR and proteomic analyses, we found that the Δ*agaSY* mutant exhibited strong upregulation of F9 fimbrial expression, accompanied by distinct fimbrial structures observed under electron microscopy. This transcriptional increase was further supported at the protein level by LC-MS/MS quantification, which revealed an increased abundance of F9 fimbrial proteins in the Δ*agaSY* mutant compared to the wild-type strain, particularly at 28°C. F9 fimbriae have been reported to mediate biofilm formation and bind Galβ1-3GlcNAc glycans (56), suggesting that their induction compensates for the loss of type 1 fimbriae to maintain adhesion. In line with these observations, similar fimbrial switching has been reported in UPEC strain CFT073 with deletion of *fim* and *pap* operons, leading to increased expression of F1C fimbriae (33). Our findings, therefore, extend the model of fimbrial hierarchy, highlighting a link between amino sugar metabolism and altered production of fimbriae. The regulatory basis for this switch remains to be defined, but may involve global regulators such as H-NS, which represses *f9* expression at 37°C (27) or metabolic regulators that respond to changes in amino sugar availability.

### Effect of *agaSY* deletion on motility

In addition to reduced fimbrial production, the Δ*agaSY* mutant displayed markedly increased swimming motility, reflecting the established crosstalk regulation between fimbriae and flagella described in UPEC (26, 32). For example, fimbrial expression is known to be decreased in favor of motility through the repression of the master flagellar regulator *flhD* by PapX, a MarR-like regulator encoded by the *pap* operon (57). Loss of type 1 fimbriae in the Δ*agaSY* mutant may relieve this repression, resulting in increased motility.

Enhanced motility may also contribute to bacterial dissemination within the urinary tract, which may help ascension from the bladder to the kidneys (26); however, excessive motility may result in decreased adherence, which represents a trade-off between colonization strategies. According to our findings, the metabolism of amino sugars may have an impact on this balance, potentially by modulating the expression of regulators that coordinate fimbrial and flagellar systems.

### Attenuation of the *agaSY* mutant in the murine UTI model

Given the multiple effects of the loss of Aga/Gam sugar metabolism caused by deletion of *agaSY* genes, we hypothesized that these genes may also be important for colonization of the urinary tract in a murine model. Our *in vivo* experiments confirmed that the Δ*agaSY* mutant is significantly attenuated for colonization of both the bladder and kidneys, while complementation restored bacterial loads to wild-type levels. These findings highlight the importance of amino sugar metabolism for UPEC fitness in the urinary tract, a challenging environment characterized by limited nutrient availability, iron sequestration, and host immune defenses (1, 6). Previous studies have established that UPEC virulence requires gluconeogenesis, the TCA cycle, and iron acquisition systems (7). Our work highlights the interaction between metabolism and virulence by adding *agaSY* to the list of metabolic genes that contribute to infection.

The attenuation observed in the Δ*agaSY* mutant is likely multifactorial. Reduced type 1 fimbriae expression decreases adhesion to bladder epithelial cells, impaired biofilm formation may limit persistence and altered fimbrial expression patterns can affect colonization dynamics. Together, these defects result in reduced bacterial fitness in the urinary tract. This is consistent with the essential role of type 1 fimbriae in preventing urine flow and establishing infection (10, 25).

Beyond the regulatory networks already described in ExPEC, studies on Aga/Gam metabolism in *Streptococcus suis* and the function of GalNAc in immune response (58) indicate that amino sugar catabolism may play a role in bacterial pathogenesis through metabolic pathways as well as regulatory pathways, in infection models in mice and piglets (31). In addition, in the non-pathogenic *E. coli* K-12, N-glucosamine (Nag) has been linked to regulation of type 1 fimbriae (59) while NagC, another gene of the Nag operon, has been implicated in host colonization in *Vibrio fischeri,* facilitating colonization of its squid host, *Euprymna scolopes* (60). Together, these findings support a model in which amino sugar metabolism intersects with fimbrial expression, motility, and biofilm formation, ultimately influencing bacterial fitness and virulence in the urinary tract.

### Broader implications of amino sugar metabolism for virulence regulation

Some mucus-derived sugars, such as Aga and Gam, can act as regulatory signals that influence bacterial colonization and adhesion. Recent studies have shown that in Enterohemorrhagic *E. coli* (EHEC), bacteria can exploit fucose, GalNAc and N-acetylneuraminic acid as signaling molecules to modulate metabolism and regulate the expression of their virulence factors (61). Similarly, in the adherent-invasive *E. coli* (AEIC) LF82 strain, the transcriptional regulator NagC has been implicated in the early steps of biofilm formation, highlighting how bacterial fitness can be influenced by sugar sensing and metabolism. In our study, we demonstrated that amino sugar metabolism, mediated by *agaSY*, not only enables UPEC to utilize alternative carbon sources but also regulates surface structures critical for virulence. Because of their dual role, metabolic pathways are attractive targets for treatments. Disruption of such pathways could simultaneously impair bacterial growth and attenuate virulence, offering a double therapeutic benefit.

The need for new treatment approaches is urgent due to the rise of multidrug-resistant UPEC strains (5). Because metabolic pathways are subject to physiological constraints, they may be less mutable targets compared with classical virulence factors. Moreover, inhibitors of amino sugar metabolism could synergize with existing antibiotics by reducing bacterial fitness and decreasing biofilm formation.

Future studies should explore the regulatory mechanisms linking amino sugar metabolism to fimbrial expression. Transcriptomic and metabolomic analyses of Δ*agaSY* mutants under infection-relevant conditions may identify global regulators or signaling molecules mediating this connection. Finally, it will be important to assess whether similar phenotypes are observed in clinical multidrug-resistant UPEC isolates, to evaluate the translational potential of targeting amino sugar pathways as a therapeutic strategy.

### Influence of *agaSY* on carbon catabolism in UPEC

The capacity of UPEC to exploit the various carbon sources in human urine is widely regarded as a key environmental adaptation. Persistence in the urinary tract depends on the ability to use different carbon sources because it is a nutrient-limited niche (1). In line with the known function of AgaS as an isomerase and AgaY as an aldolase in the Aga/Gam metabolic pathway (14, 17), we demonstrate in this study that deleting *agaSY* in UPEC strain CFT073 results in an incapacity to utilize Aga as the sole carbon source. Interestingly, despite the fact that the CFT073 genome contains the complete *aga/gam* gene clusters, including the *agaBCDI* genes required for Gam transport, it does not grow on Gam. This is in contrast to *E. coli* W, which can grow on Gam or Aga as sole carbon sources. This suggests that *aga* genes may be partially inactive or regulated differently in some *E. coli* lineages. Previous studies in *E. coli* K-12 and *E. coli* O157:H7 identified natural polymorphisms in the *aga/gam* cluster that impair Gam utilization (14, 16), and our data are in line with such variability in UPEC. In addition, deletion of *agaS* and *agaY* in the UPEC strain UTI89 and the avian pathogenic *E. coli* (APEC) strain CH138 also resulted in the loss of Aga metabolism (Table S1), highlighting the conserved role of these enzymes within ExPEC for Aga utilization. Previous studies identified *agaR* as a repressor of the *aga/gam* genes under certain conditions; however, our results suggest that *agaSY* may have additional functions in pathway activity or regulation beyond this control. Transcriptomic evidence demonstrating the upregulation of *dsdA* (encoding a D-serine deaminase) in urine further supports the significance of nutrient flexibility in UPEC (8, 37). Collectively, these findings raise the possibility that similar mutations or regulatory mechanisms shape the metabolic versatility of UPEC in the urinary tract.

In addition to Aga, our phenotypic microarray assays revealed that the Δ*agaSY* mutant also displayed impaired utilization of other amino sugars, including N-acetylglucosamine (GalNAc), which is one of the most abundant amino sugars in nature and an essential precursor for peptidoglycan and lipopolysaccharide (LPS) synthesis (17). Further, the *agaSY* mutant was also unable to effectively use many other carbon sources and amino acids, as shown by the phenotypic array analysis. To confirm selected phenotypes identified in the phenotypic microarray analysis, growth kinetics were assessed in minimal medium supplemented with 02.% fructose, 0.2% galactose, or 0.2% mannose as the sole carbon source. Consistent with the phenotypic microarray data, the Δ*agaSY* mutant exhibited growth comparable to that of the CFT073 wild-type strain in the presence of fructose or mannose (Fig.S2A-B). In contrast, growth of the mutant was markedly impaired with galactose as the sole carbon source (Fig.S2C). Interestingly, transcriptomic analyses, combined with comparative evaluation of UPEC fitness in urine, have shown that the utilization of short peptides and amino acids as carbon sources plays a critical role during infection of the bladder and kidneys. Consistently, mutants deficient in peptide import exhibit reduced fitness during UTI, suggesting the importance of carbon and peptide import for UPEC survival and virulence (37, 62).

This finding highlights the central role of *agaSY* in broader amino sugar and carbon metabolism. Since amino sugars are key structural components of bacterial cell envelopes, defects in their utilization may also change the integrity of the envelope and influence other virulence-associated traits. Indeed, bacterial resistance to osmotic stress and antimicrobial peptides, both of which are present in the urinary tract, is maintained in large part by cell wall precursors made from amino sugars. In line with this, we hypothesize that amino sugar catabolism not only supports structural maintenance but may also contribute to pathogen-host interactions. Furthermore, the regulation of Aga/Gam catabolism is likely involved in the successful establishment of UPEC infection.

### Conclusions

In summary, our study shows that *agaSY* and the Aga/Gam metabolic pathway is a central metabolic determinants in UPEC, required for amino sugar utilization, and that this pathway contributes to fimbrial regulation, biofilm formation, motility, and successful colonization of the urinary tract. By investigating the effects of loss of the Aga/Gam metabolic pathway and demonstrating that nutrient availability and carbon metabolism are tightly linked to pathogenic potential, these findings offer fresh insight into the connection between metabolism and virulence and provide new perspectives on therapeutic targets against multidrug-resistant UTI pathogens.

## Materials and Methods

### Animal ethics statements

All animals used in this study were clinically healthy and had been acclimated for 7 days before experimentation. All procedures were conducted in strict accordance with the ethical guidelines approved by the INRS Institutional Animal Care Committee (CIPA) and by the rules and standards of the Canadian Council on Animal Care (2410–03).

### Bacterial strains, growth conditions and plasmids

*E. coli* strains and plasmids used in this study are listed in Table 2. *E. coli* strain CFT073 (O6:K2:H1) was isolated from the blood of a patient with acute pyelonephritis in the late 1980s (18). Bacteria were preserved at -80°C ± 10°C as glycerol stocks and cultured in Lysogeny Broth (LB) (Alpha Bioscience, Baltimore, MD) at 37°C following one or more passages on LB agar plates to ensure culture purity. When required, antibiotics and reagents were added at the following final concentrations: kanamycin (50 μg/mL), ampicillin (100 μg/mL), and chloramphenicol (30 μg/mL).

**Table 2.** Bacterial strains and plasmids used in this study.

| Bacterial strains | Relevant genotype | References |
| --- | --- | --- |
| CFT073 | UPEC wild-type pyelonephritis strain (O6:K2:H1) | (18) |
| QT775 | χ7213 (MGN-617) *thi thr leu tonA lacY glnV supE ΔasdA4 recA::RP4 2-Tc::Mu [λpir]*, Km<sup>r</sup> | (19) |
| QT5158 | CFT073 *ΔagaS::FRT* | This study |
| QT6076 | CFT073 *ΔagaY::FRT* | This study |
| QT6077 | CFT073 *ΔagaSY::FRT* | This study |
| QT6105 | CFT073 *attTn7-Cm::agaSY*, Cm<sup>r</sup> | (11) |
| QT2138 | CFT073 strain *Δfim::Km<sup>r</sup>* | (20) |
| QT2285 | CFT073 *fimS* phase L-ON, Cm<sup>r</sup> | (20) |
| QT3034 | *E. coli* W | ATCC 9637 |
| QT2799 | *Serratia liquefaciens* | ATCC 27592 |
| QT4652 | *E. coli* K-12 BW25113 | (21) |
| Plasmids | Relevant genotype | References |
| pKD4 | Template plasmid for the amplification of the *km* cassette bordered by FRT sites | (21) |
| pKD46 | λ-Red recombinase plasmid Ts replicon; Ap<sup>r</sup> | (21) |
| pIJ258 (pSTNSK) | pST76-K::*tnsABCD*, Km<sup>r</sup> | (19) |
| pIJ360 (pSTNSK) | pGP-Tn7-FRT:: Cm, Ap<sup>r</sup>, Cm<sup>r</sup> | (19) |
| pCP20 | FLP helper plasmid Ts replicon; Ap<sup>r</sup> Cm<sup>r</sup> | (21) |
| pGP-Tn7-Cm | pGP-Tn7-FRT:: Cm, Ap<sup>r</sup>, Cm<sup>r</sup> | (19) |
| pIJ628 | pGP-Tn7-Cm::*agaSY*; Ap<sup>r</sup>, Cm<sup>r</sup> | This study |

### Construction of site-directed mutants and complementation of strains

All mutant strains were generated using the method described by Datsenko and Wanner, with plasmid pKD4 serving as the template for amplification of the kanamycin resistance cassette (21). The primers used for gene disruption are listed in Table 2. In the resulting mutant strains, antibiotic resistance cassettes flanked by FLP recombination target (FRT) sites were excised by introducing pCP20, which expresses the FLP recombinase (22).

The mutant strains were complemented by inserting the respective genes at the *att*Tn*7* chromosomal site. This procedure was carried out as described by S. Crépin *et al.*(19). Briefly, the *agaS* and *agaY* genes and their native promoters were amplified using primers CMD2322 and CMD2990 as listed in Table 2. The PCR product was digested with *XhoI* and *KpnI* and subsequently ligated into the mini-Tn*7* vector pGP-Tn*7*-Cm. Conjugation between the donor strain MGN-617 (χ7213) carrying the pGP-Tn*7*-*agaSY* and the CFT073 Δ*agaSY* recipient strain containing the helper plasmid pSTNSK (*tnsABCD*) was done at 30°C ± 2°C on LB agar plates supplemented with diaminopimelic acid (DAP). The recipient mutant containing the genes for complementation was selected by streaking on LB agar containing 30 μg/mL of chloramphenicol and incubating overnight at 37°C ± 2°C. The resulting colonies were screened with primers CMD1298 and CMD1421 (Table 3) to confirm the successful site-specific integration of the Tn*7*-*agaSY* construction at the *att*Tn*7* locus.

**Table 3.**
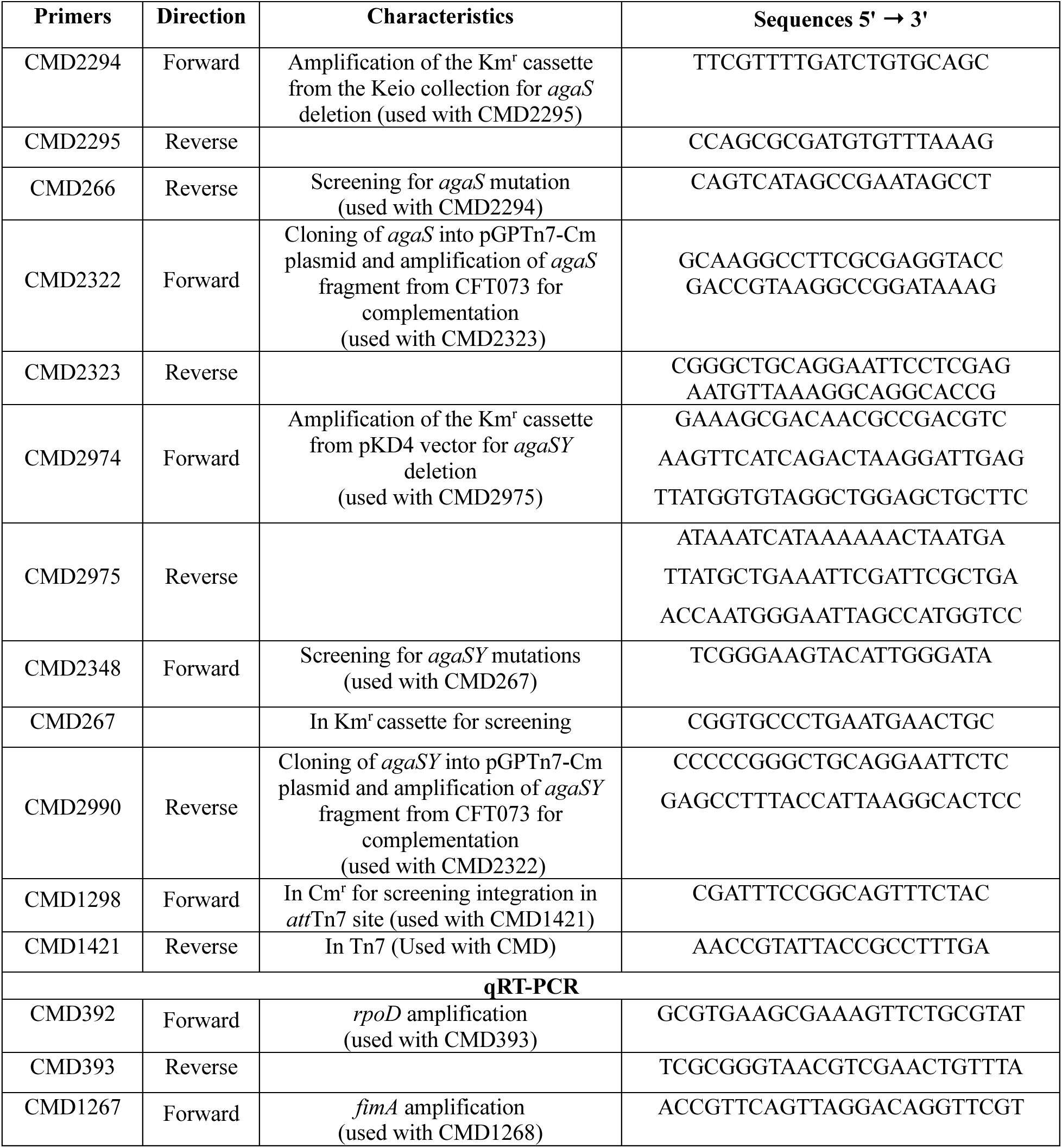
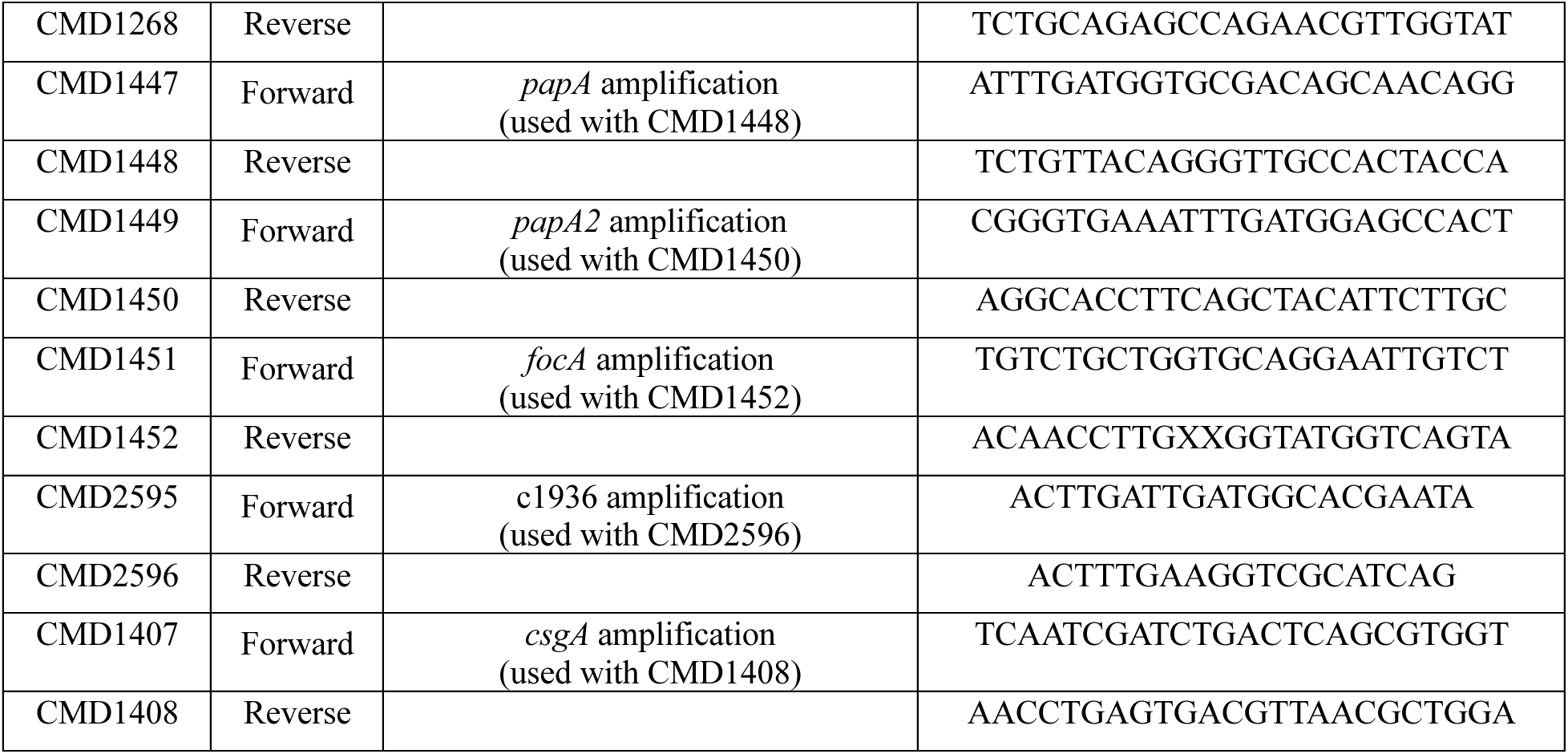
PCR and qPCR primers used in this study.

### Sugar utilization assays

The CFT073 wild-type strain, its derivative mutant, CFT073 Δ*agaSY*, its complemented strain CFT073 cp *agaSY*, as well as *E. coli* W (positive control) and *E. coli* K-12 (negative control), were grown overnight in 5 ml of LB broth at 37°C with shaking. Following incubation, bacterial cultures were centrifuged and washed with phosphate-buffered saline (PBS 1X, pH 7.4). For sugar utilization assays on solid media, bacteria were grown on M9 minimal agar plates supplemented with one of the following sole carbon sources (20 mM): N-acetyl-D-galactosamine (Aga), D-galactosamine (Gam), glucose, xylose, or galactose. The M9 medium contained essential components for bacterial growth: For a final concentration in 1L, i) a source of nitrogen and phosphorus (Na₂HPO₄·7H₂O, 0.04 M; KH₂PO₄, 0.02 M); ii) sodium (NaCl, 0.02 M); iii) calcium (CaCl₂, 1 mM); iv) magnesium (MgSO₄·7H₂O, 1 mM); v) thiamine (0.5 mM); and vi) the tested carbon source (20 mM). Plates were incubated at 37°C, and bacterial growth was assessed after at least 48 hours.

For liquid-phase assays, kinetic growth was monitored using the automated turbidimetric Bioscreen C system (Oy Growth Curves Ab Ltd., Raisio, Finland). Briefly, 2 μL of overnight culture was inoculated into 200 μL of M9 minimal broth supplemented with either 20 mM glucose, Aga, or Gam. Microtiter plates were incubated at 37°C, with optical density (OD_600_) measurements recorded every 15 minutes over 48 h. Before each measurement, the plates were agitated for 10 seconds at medium amplitude. Each condition was tested in ten replicate wells per strain.

### Phenotypic microarrays

Metabolic profiles of CFT073 wild-type strain and its derivative mutant CFT073 Δ*agaSY* were determined using the phenotypic microarray (PM) Omnilog™ system (Hayward, CA, USA), which can measure and compare growth of different strains upon exposure to hundreds of growth conditions over 48 hours. Experiments were conducted following the Omnilog™ manufacturer’s protocol with some modifications (23). All materials, media, PM and reagents for the PM system were purchased from Omnilog™. Briefly, bacterial strains were grown on an LB agar plate at 37°C for 16-18 hours. Then, using a sterile inoculum loop, colonies were resuspended in 15 mL of IF-0 minimal medium. Bacterial density was adjusted to 42% transmittance (T) using a turbidimeter. These suspensions were subsequently diluted sixfold by adding 75 mL of IF-0 medium to reach a final density of 85% T. In this study, a PM1 96-well plate containing different carbon sources was used to assess the metabolic utilization of various substrates. The plates were inoculated with 100 μL of either CFT073 wild-type or the CFT073 Δ*agaSY* mutant and incubated at 37°C in an OmniLog™ device. Color development, indicative of bacterial respiration, was monitored for 48 hours. The reduction of tetrazolium violet (dye A) within each well served as a reporter, and color intensity was measured in arbitrary units (AU), with a maximum of 500 AU. Kinetic data were analyzed using OmniLog-PM software from Biolog, and all assays were performed in duplicate for each bacterial strain.

### Evaluation of type 1 fimbriae production mediated by yeast agglutination and Pap fimbriae by hemagglutination

The capacity of bacterial strains to produce type 1 fimbriae was measured by the ability to agglutinate *Saccharomyces cerevisiae* cells on a polystyrene surface, adapted from(24). Briefly, strains were cultured in LB broth at 37°C until reaching OD_600_ 0.6 ± 0.02 with shaking at 250 rpm. Bacterial cells were harvested and concentrated in PBS to a final suspension of approximately 2 × 10^11^ cells/mL. Subsequently, 40µL of each diluted bacterial suspension was added to a round-bottom 96-well plate and serially diluted 2-fold. An equal volume of 3% (v/v) commercial yeast suspension was added to each well. After 30 minutes on ice, yeast aggregation was assessed. Agglutination was monitored visually by aggregation and precipitation of the cells. The agglutination titer was defined as the highest dilution at which visible yeast aggregation occurred. An isogenic CFT073 *fim* knockout mutant (Δ*fim*) was used as a negative control, while a CFT073 *fim*-locked-ON mutant served as a positive control.

The presence of P fimbriae was demonstrated by the mannose-resistant hemagglutination (MRHA) of human erythrocytes (blood type O+, from volunteer donors). Under the same conditions as those used for yeast agglutination, bacterial strains were cultured to mid-log phase. Subsequently, 50 μL of a 3% (v/v) suspension of human red blood cells was added to each well of a round-bottom 96-well plate containing the bacterial suspension. To inhibit type 1 fimbriae-mediated hemagglutination, α-D-mannopyranose was added to a final concentration of 2.5%.

### Adhesion assay

Adhesion of bacteria to the bladder epithelium is achieved through the production of type 1 fimbriae, which are essential for the establishment of UPEC infection. The urothelial cell adhesion test is used to verify whether deletion of the gene encoding the *agaSY* genes can affect the virulence of UPEC strain CFT073 by improving or decreasing adherence to bladder epithelial cells. The established test is based on experiments by Martinez (2000)(25). Human bladder 5637 cells (ATCC HTB-9) were cultured in RPMI medium (Wisent Bioproducts) supplemented with 10% fetal calf serum, 2 mM L-glutamine, 10 mM HEPES, 1 mM sodium pyruvate, and 4.5 g/L glucose. For analysis, cells were grown to confluence in RPMI, and 5 × 10^5^ cells/well were distributed in 24-well plates. UPEC strain CFT073 and the isogenic *agaSY* mutant were grown in LB medium at 37°C to an OD_600_≈0.6 ± 0.02. Before infection, cultures were washed with 1 x PBS, suspended in RPMI medium and inoculated at a multiplicity of infection (MOI) of 10 CFU per epithelial cell. Contact between bacteria and epithelial cells was increased by centrifugation at 500 x *g* for 6 min. After 2 hours, to collect only adherent bacteria, cells were washed three times with PBS and lysed with PBS-sodium deoxycholate (DOC) to a final concentration of 0.1%. Then, each bacterial suspension was collected, serially diluted and spread on MacConkey agar plates to quantify bacteria.

### Motility experiments

Swimming motility experiments were performed as previously described by Lane, Lockatell *et al.*(26), with the following modifications. First, bacterial strains were grown until mid-log exponential phase (OD_600_ 0.6 ± 0.02) in LB broth at 37°C with shaking. Next, with a sterile inoculating loop, cultures were stabbed into the center of either soft agar plates (1% tryptone, 0.5% NaCl, 0.25% agar) or soft agar minimal M9 medium plates, containing either 20mM glucose, 20 mM Aga, or 20 mM Gam as a carbon source and 0.25% pure agar. Plates were prepared one day before inoculation and left to solidify at room temperature overnight. During inoculation, care was taken to avoid contact with the bottom to prevent twitching motility. Inoculated plates were incubated at 37°C for 16 to 22 hours for soft agar assays, and an extended period of 48 to 72 hours when using minimal soft agar media. Bacterial motility was assessed by measuring the diameter of the migration zone in millimeters (mm). All experiments were performed in triplicate.

### Bacterial RNA extraction and quantification of fimbrial gene expression

Bacterial cultures were grown in triplicate in LB broth under the same conditions as used for the adhesion assays. Once the OD_600_ reached approximately 0.6, 500 µL of each culture was collected into 1.5 mL Eppendorf tubes, and 1 mL of Bacterial RNAprotect (Qiagen, Toronto, ON, Canada) was added to the cultures. They were then processed following the protocol. Bacterial total RNA was extracted using the RNeasy Mini Kit (Qiagen, Hilden, Germany, Cat. No. 74104), following the protocol recommended by the manufacturer. Genomic DNA contamination was removed using the Turbo DNA-free™ Kit (Invitrogen, Thermo Fisher Scientific, Waltham, MA, USA) according to the manufacturer’s protocol. RNA concentration was determined using a Nanodrop (ND-1000), and RNA integrity was assessed by agarose gel electrophoresis, with visualization of the 16S and 23S RNA bands. Complementary DNA (cDNA) was synthesized using the Wisent Advanced™ cDNA Synthesis Kit (Wisent Bioproducts, St-Bruno, Québec, Canada) according to the manufacturer’s instructions. Quantitative real-time PCR (qRT-PCR) was then performed using Wisent Advanced™ qPCR Mastermix with Supergreen™ dye (Wisent Bioproducts, St-Bruno, Québec, Canada) following the manufacturer’s instructions. The *rpoD* gene served as an internal housekeeping control. Each qRT-PCR reaction was carried out in triplicate. The threshold cycle (Ct) values were calculated and normalized to the corresponding reference gene, *rpoD* Ct values, and relative gene expression levels were calculated using the 2^−ΔΔCt^ method, followed by log₂. Genes with a fold-change greater than or less than the defined threshold of 1 were considered differentially expressed. Primers used for the qRT-PCR analysis are listed in **Error! Reference source not found.**.

### Electron microscopy

Bacterial cells were grown under the same conditions as those used for the agglutination assay. A glow-discharged Formvar-coated copper grid was placed on a drop of bacterial suspension and incubated for 15 minutes to allow cell adsorption. Excess liquid was carefully removed with filter paper, followed by the application of a drop of 1% phosphotungstic acid (negative stain) for 15 seconds. Grids were then air-dried and examined using a Hitachi H-7100 transmission electron microscope (Toronto, Ontario, Canada).

### Preparation of fimbrial extracts and quantification

Fimbrial extracts were prepared as described previously (27). Briefly, bacteria were cultured in 50 mL of LB with 100 µL of an overnight preculture and incubated for 16-18 hours at 37°C or 28°C. Cells were harvested at 10,000 x *g* for 10 min at 4°C and washed once with 25 mL of PBS at 4°C. The bacterial pellet was resuspended in 1 mL of EDTA buffer (pH 7.4) and incubated 30 min at 56°C. Subsequently, cells were centrifuged at 10,000 x *g* for 10 min at 4°C to collect the supernatant containing fimbrial proteins. Then, proteins were precipitated by adding trichloroacetic acid (TCA) to a final concentration of 20% (w/v) and incubated overnight at 4°C. Proteins were washed once with Tris-EDTA (TE) buffer (pH 12) and twice with TE buffer (pH 8), then resuspended in 500 µL of TE buffer pH 8. Protein samples were digested with trypsin and analyzed by liquid chromatography-tandem mass spectrometry (LC-MS/MS) at the Proteomic Core Facility, Center for Advanced Proteomic Analyzes (CAPA) of the Institute for Research in Immunology and Cancer (IRIC, Université de Montréal, Montréal, Québec, Canada) for protein identification and quantification.

### Biofilm formation

Biofilm formation on polystyrene surfaces was conducted in 96-well plates as detailed in earlier studies (28). Bacterial strains were grown in LB medium at 37°C ± 2°C until the exponential phase. Once the bacterial cultures reached an OD_600_≈0.6, 200 μL of each suspension was dispensed into each well of a 96-well U-bottom plate. The plates were incubated statically at 37°C for 48 hours to allow biofilm formation. After incubation, the wells were washed twice with PBS to remove non-adherent cells, then stained using 0.1% crystal violet solution for 30 minutes at 37°C. Excess stain was removed by washing the wells four times with PBS 1X. The plates were then air-dried, and biofilm-bound dye was solubilized by adding a solution of acetone and ethanol (80:20 v/v) to each well. The biofilm density was quantified by measuring the absorbance at OD_595nm_ using a microplate reader.

### Experimental UTI in CBA/J mice

A murine ascending UTI model was used, where experimental infections were carried out as described previously(29). Prior to inoculation, the strains were grown for 16 hours at 37°C with shaking (250 rpm) in 55 ml of LB medium. For each group 10 mice were infected with 30 μL of a pure culture (ranging from 1.10 to 2.40 × 10^9^ CFU/30 μL) with either the CFT073 wild-type strain, the mutant CFT073 Δ*agaSY* or its complemented derivative strain CFT073 cp *agaSY*. At 48 hours post-infection (p.i.), mice were euthanized, and bladders and kidneys were aseptically harvested. Then, the organs were weighed, homogenized, serially diluted in 96-well plates and subsequently plated on MacConkey agar plates to quantify bacterial counts.

### Statistical analysis

All statistical analyses were performed using GraphPad Prism 10 software (GraphPad Software, San Diego, CA, USA). Statistically significant differences between two groups were established by an unpaired t-test, and comparisons among three or more groups were performed by the Kruskal-Wallis test, followed by Dunn’s multiple comparisons test to compare the wild-type, the mutant and the complemented strain. Statistical significance was defined as * *p≤* 0.05; ** *p≤* 0.01; *** *p≤* 0.001; and **** *p≤* 0.0001.

## Acknowledgments

We thank all the collaborators and colleagues who helped in this study. This research was funded by the Natural Sciences and Engineering Research Council (NSERC) Canada: Grant Number RGPIN-2025-07045, the Canadian Institutes of Health Research Grant Number FRN168006 and the Genome Québec special AMR program.

## Author Contributions

Carole Anamalé, Hicham Bessaiah and Charles M. Dozois conceptualized the research and analyzed the data. Carole Anamalé and Hicham Bessaiah performed the experiments and wrote the manuscript. Hicham Bessaiah and Charles M. Dozois reviewed and edited the manuscript. Carole Anamalé, Hicham Bessaiah and Sébastien Houle performed the *in vivo* experiments. We thank Roger Levesque’s group, including Sima Mohammadi, for helping with Biolog experiments. We also thank Maëva Legros for her help in constructing the complemented strain and performing the assays in M9 medium, and Enya Koroghli for the adhesion assays and qRT-PCR experiments. All authors revised the manuscript critically and approved the submission of the manuscript.

## Data availability statement

All data relating to this article are present in the article and the accompanying Supplemental Information.

## Supporting Information

